# Heat wave event facilitates defensive responses in invasive C3 plant Ambrosia artemisiifolia L. under elevated CO_2_ concentration to the detriment of insect herbivores

**DOI:** 10.1101/2021.03.13.435250

**Authors:** Zhenya Tian, Chao Ma, Chenchen Zhao, Yan Zhang, Xuyuan Gao, Zhenqi Tian, Hongsong Chen, Jianying Guo, Zhongshi Zhou

## Abstract

To predict and mitigate the effects of climate change on communities and ecosystems, the joint effects of extreme climatic events on species interactions need to be understood. Using the common ragweed (*Ambrosia artemisiifolia* L.)—leaf beetle (*Ophraella communa*) system, we investigated the effects of heat wave and elevated CO_2_ on common ragweed growth, secondary metabolism, and the consequent impacts on the beetle. The results showed that elevated CO_2_ and heat wave facilitated *A. Artemisiifolia* growth; further, *A. artemisiifolia* accumulated large amounts of defensive secondary metabolites. Being fed on *A. artemisiifolia* grown under elevated CO_2_ and heat wave conditions resulted in the poor performance of *O. communa* (high mortality, long development period, and low reproduction). Overall, under elevated CO_2_, heat wave improved the defensive ability of *A. artemisiifolia* against herbivores. This implies that heat wave event will relieve harm of *A. artemisiifolia* to human under elevated CO_2_. On the other hand, super adaptability to climatic changes may aggravate invasive plant distribution, posing a challenge to the control of invasive plants in the future.

## Introduction

Global warming is one of the hottest issues and has the potential to dramatically change environments worldwide (Vinagre et al., 2018). It is known that climate change is likely to impact species both directly and indirectly (Sun et al., 2018; Tong et al., 2017; Drake et al., 2017). Heat wave events are a universal climatic phenomenon in meteorology and occur when the daily maximum temperature of more than five consecutive days exceeds the average maximum temperature by 5°C (García-Herrera et al., 2010). In recent years, heat waves have imposed severe stresses on natural and human systems (Seo et al., 2019; Skinner et al., 2018). Heat waves can increase crop failure and the loss of livestock (García-Herrera et al., 2010) and reduce the gross primary productivity of vegetation (Ciais et al., 2005). Heat waves will increase in intensity, duration, and frequency because of global climate change (Seo et al., 2019). On the other hand, increasing CO_2_ is also a common phenomenon in global climate change (Sun et al., 2018). In 2016, the CO_2_ concentration was 46% higher than that in pre-industrial times (WMO, 2018), and it is predicted to reach 750 ppm by the end of the century (IPCC, 2013; Meehl et al., 2006). Elevated CO_2_ is known to alter an array of plant traits, and the typical effects of elevated CO_2_ on plants include increases in the photosynthetic rate, biomass, and carbon: nitrogen (C: N) ratio (Ainsworth &Rogers, 2007; Korner, 2000). In addition, elevated CO_2_ can regulate the assimilation and allocation of C and N resources within plant tissues and alter the primary and secondary metabolites of plants (Tong et al., 2018). Many previous studies have demonstrated that, under elevated CO_2_ conditions, secondary defensive substances increase in plants, thus indirectly affecting the development and reproduction of herbivorous insects (Schädler et al., 2007; Curtis & Wang, 1998; Poorter et al., 1997; Lincoln et al., 1993). However, even more dramatic reductions in the larval performance of herbivorous insects have been observed in some herbaceous C3 plants under elevated CO_2_ conditions (Karowe et al., 2007). Previous studies have revealed that mortality over the entire larval period increased by 180% for buckeye larvae feeding on plantain grown at 700 ppm (Fajer et al., 1989) and by 135% for leaf miners feeding on Paterson’s Curse (Johns & Hughes, 2002).

It is known that higher temperatures are always accompanied by elevated atmospheric CO_2_ concentrations (Lee et al., 2020), and the global mean surface temperature is expected to rise by 1–4.5°C, depending on the scenario, by the year 2100 (IPCC, 2013). Few case studies have explored the effects of elevated CO_2_ and elevated temperature on herbivore performance in such a way that both individual and combined effects of these two factors could be evaluated (Zvereva & Kozlov, 2006). Several studies have concluded that temperature does not affect plant and herbivorous insect responses to CO_2_ elevation (Williams et al., 2000; Buse et al., 1998), while others have demonstrated strong interactive effects of CO_2_ and temperature (Johns et al., 2003; Johns & Hughes, 2002; Veteli et al., 2002). However, a more recent study has demonstrated that relatively low concentrations of plant secondary metabolites were observed when plants were exposed to the combined impact of elevated CO_2_ and temperature, and the responses of both phenolics and terpenes to CO_2_ were strongly modified by elevated temperatures. Therefore, elevated temperatures can lead to the amelioration of the negative effects of CO_2_ on herbivore fitness (Zvereva & Kozlov, 2006);in fact, elevated temperatures may counteract the defense-inducing effect of elevated CO_2_ on plants (Veteli et al., 2007; Zvereva & Kozlov,2006;Veteliet al., 2002).However, even when the total heat sum is the same, compared with a long-term increase in constant temperature, heat waves have more severe negative effects on plant physiology and communities (Bauweraerts et al., 2014).

Common ragweed, *Ambrosia artemisiifolia* L. (Asteraceae) is a noxious and invasive weed originating from North America. It is a C3 plant that often causes large losses to agriculture (Cowbrough & Brown, 2003). In addition, 13.5 million persons suffer from *Ambrosia*-induced allergies in Europe, causing costs of €7.4 billion annually (Schaffner et al., 2020). Some previous studies have revealed that the growth and reproduction of *A. artemisiifolia* were enhanced when grown under elevated CO_2_ conditions (Stinson & Bazzaz, 2006), and its pollen production was also dramatically increased at elevated CO_2_ levels (Rogers et al., 2006; Wayne et al., 2002; Ziska & Caulfield, 2000). Physiological and morphological mechanisms by which elevated CO_2_ enhances the status of this species as an agricultural pest and allergenic weed have been demonstrated (Stinson et al., 2006). Increased levels of individual flavonoid metabolites in *A. artemisiifolia* were found under elevated CO_2_ conditions (Kelish et al., 2014). Currently, however, there are no reports on the impact of climate change on common ragweed’s natural herbivorous enemies, which is crucial for the prediction of common ragweed–herbivorous interactions in the future.

In this study, we first examined whether the secondary metabolites of the C3 plant *A. artemisiifolia* increase when the plant experiences heat wave events at elevated CO_2_ levels. Second, we explored whether secondary metabolic changes in *A. artemisiifolia* have adverse effects on the performance of the specialist herbivorous insect *O. communa*. Based on the results of this study, we will be able to predict whether exposure to heat wave events under elevated CO_2_ conditions in the future will facilitate the defensive ability of these invasive C3 plants to withstand insect herbivores more effectively. In addition, we will be able to predict whether heat wave event will relieve harm of *A. artemisiifolia* to human under elevated CO_2_ in the future.

## Results

### Effects of high CO_2_ concentration and heat wave on the performance and foliar chemistry of common ragweed

#### Leaf photosynthesis

In both temperature treatments, leaf photosynthesis in *A. artemisiifolia* under elevated CO_2_ was significantly higher than that under ambient CO_2_ during this study (1^st^ measurement: *F*_3,36_ = 219.32, *P*< 0.0001; 2^nd^: *F*_3,36_ = 162.97, *P*< 0.0001; 3^rd^: *F*_3,36_ = 122.49, *P*< 0.0001; 4^th^: *F*_3,36_ = 189.04, *P*< 0.0001; and 5^th^: *F*_3,36_ = 160.19, *P*< 0.0001). Under elevated CO_2_, leaf photosynthesis at the ambient air temperature was significantly higher than that in the heat wave treatment in all measurements except for the fourth measurement time period (day 15 post-heat stress). Meanwhile, there was no significant difference in leaf photosynthesis rate between ambient temperature and heat wave treatments under ambient CO_2_ conditions (*P* > 0.05), except for the 5^th^ measurement, i.e., on day 20 post-heat stress (Supplementary Fig. 1).

#### Total leaf area

The total leaf area of *A. artemisiifolia* under elevated CO_2_ was significantly higher than that under ambient CO_2_ in both the temperature treatments (ambient and heat wave) (*F*_3,16_ = 5.04, *P* = 0.012; Supplementary Fig. 2). Under elevated CO_2_conditions, the total leaf area of *A. artemisiifolia* was significantly increased in the ambient temperature treatment compared with the heat wave stress treatment (*P*< 0.05). In addition, under ambient CO_2_, the total leaf area at ambient temperature was higher than that in the heat wave treatment, although these differences were not significant (*P*> 0.05).

#### Height

At all five post-heat stress measurement periods, the heights of *A. artemisiifolia* under elevated CO_2_ were significantly greater than those under ambient CO_2_ (1^st^ measurement: *F*_3,16_ = 7.39, *P* = 0.0025; 2^nd^: *F*_3,16_ = 5.29, *P*< 0.0100; 3^rd^: *F*_3,16_ = 5.15, *P*< 0.0111; 4^th^: *F*_3,16_ = 3.74, *P*< 0.0329; and 5^th^: *F*_3,16_ = 4.82, *P*< 0.0141; Supplementary Fig. 3). However, there was no significant difference in plant height between ambient temperature and the heat wave stress treatment under either ambient or elevated CO_2_ (*P*> 0.05).

#### Total phenolics

The combined treatment of elevated atmospheric CO_2_ concentration and heat wave (EC+HW) had a strong effect on the total phenolic contents in *A. artemisiifolia* leaves. Throughout the experiment, the highest concentration of total foliar phenolics was in the EC+HW treatment, followed by the heat wave treatment (AC+HW); the lowest total phenolics occurred under the ambient atmospheric CO_2_ concentration and ambient temperature treatment (AC+AT) (Fig.1). At all five sampling times, *A. artemisiifolia* treated with EC+HW had a significantly higher total phenolic concentration than that in *A. artemisiifolia* leaves in the other three treatments. Except at the third sampling time, heat wave remarkably increased the total phenolic concentration at both concentrations of CO_2_ (elevated and ambient). In both the ambient temperature and heat wave treatments, elevated atmospheric CO_2_ concentration also increased total phenolics concentration on the 1^st^, 2^nd^and 4^th^ sampling times (1^st^:*P* = 0.001; 2^nd^: *P* = 0.000; 3^rd^: *P* = 0.002; 4^th^: *P* = 0.001; and 5^th^: *P* = 0.002).

**Fig.1.**
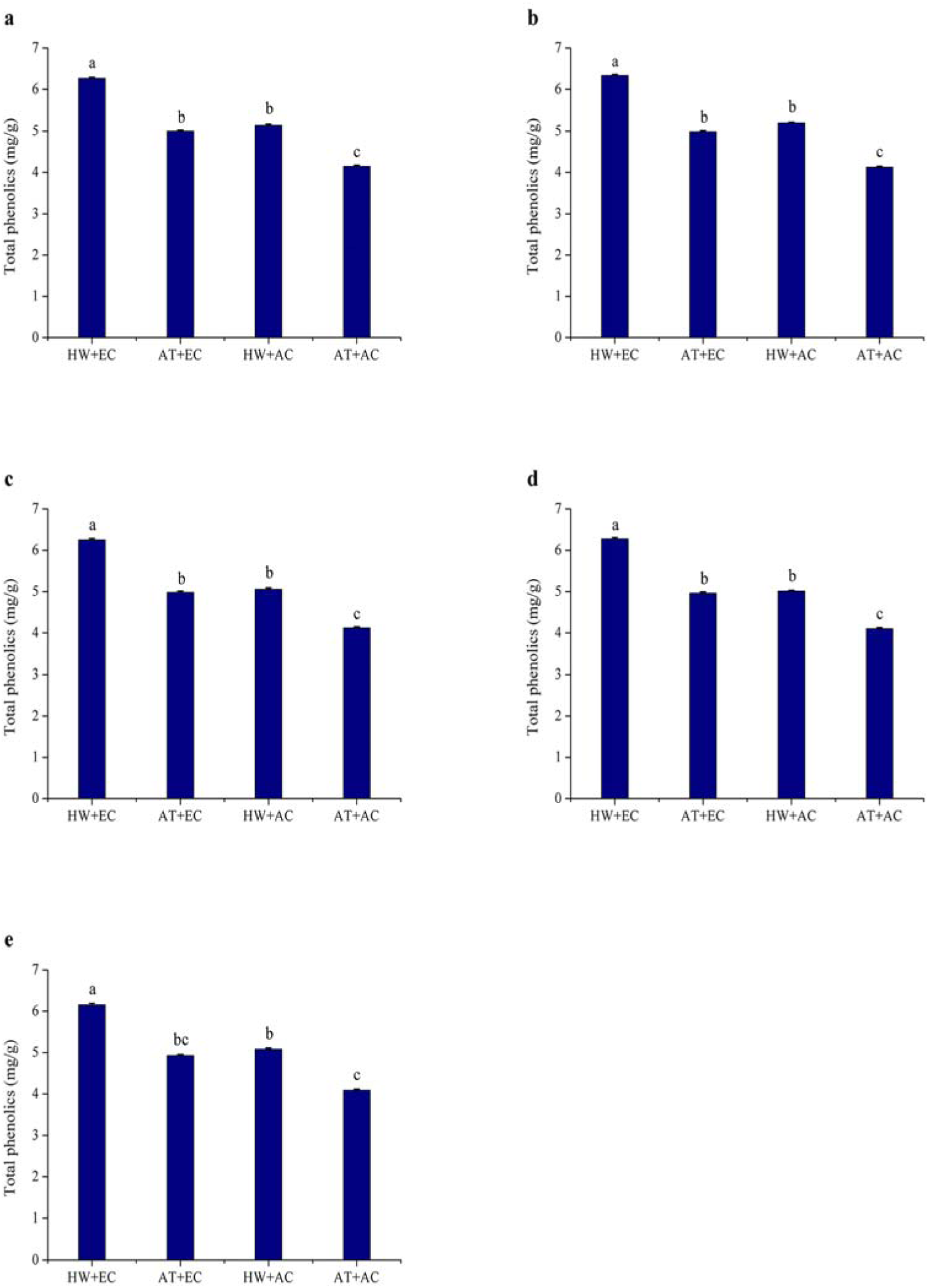
Total phenolic concentration (±SE) of *Ambrosia artemisiifolia* under different stress conditions. Data represented by columns bearing the same letters were not significantly different (LSD, *P* = 0.05). A, B, C, D, and E represent the 1^st^, 2^nd^, 3^rd^, 4^th^, and 5^th^ measurements, respectively. AT denotes the normal temperature condition; HW denotes the heat wave condition. EC and AC represent elevated atmosphere CO_2_ concentration and ambient atmosphere CO_2_ concentration, respectively.

#### Condensed tannins

Compared with the untreated control (AC+AT), elevated atmospheric CO_2_ concentration (EC+AT), heat wave (AC+HW), and their combination (EC+HW) significantly increased condensed tannin concentration in leaves at all five sampling times. Foliar condensed tannin concentrations were the highest in the EC+HW treatment, followed by the AC+HW treatment and finally the AC+AT treatment. Concentrations of condensed tannins were the lowest in AC+AT at all five sampling times (Fig.2). Regardless of CO_2_ concentration (or temperature), heat wave (or elevated atmospheric CO_2_ concentration) had a strongly positive effect on total tannin concentration in *A. artemisiifolia* leaves throughout the experiment (1^st^: *P <*0.001; 2^nd^: *P <*0.001; 3^rd^: *P <*0.001; 4^th^: *P <*0.001; and 5^th^: *P*<0.001).

**Fig. 2.**
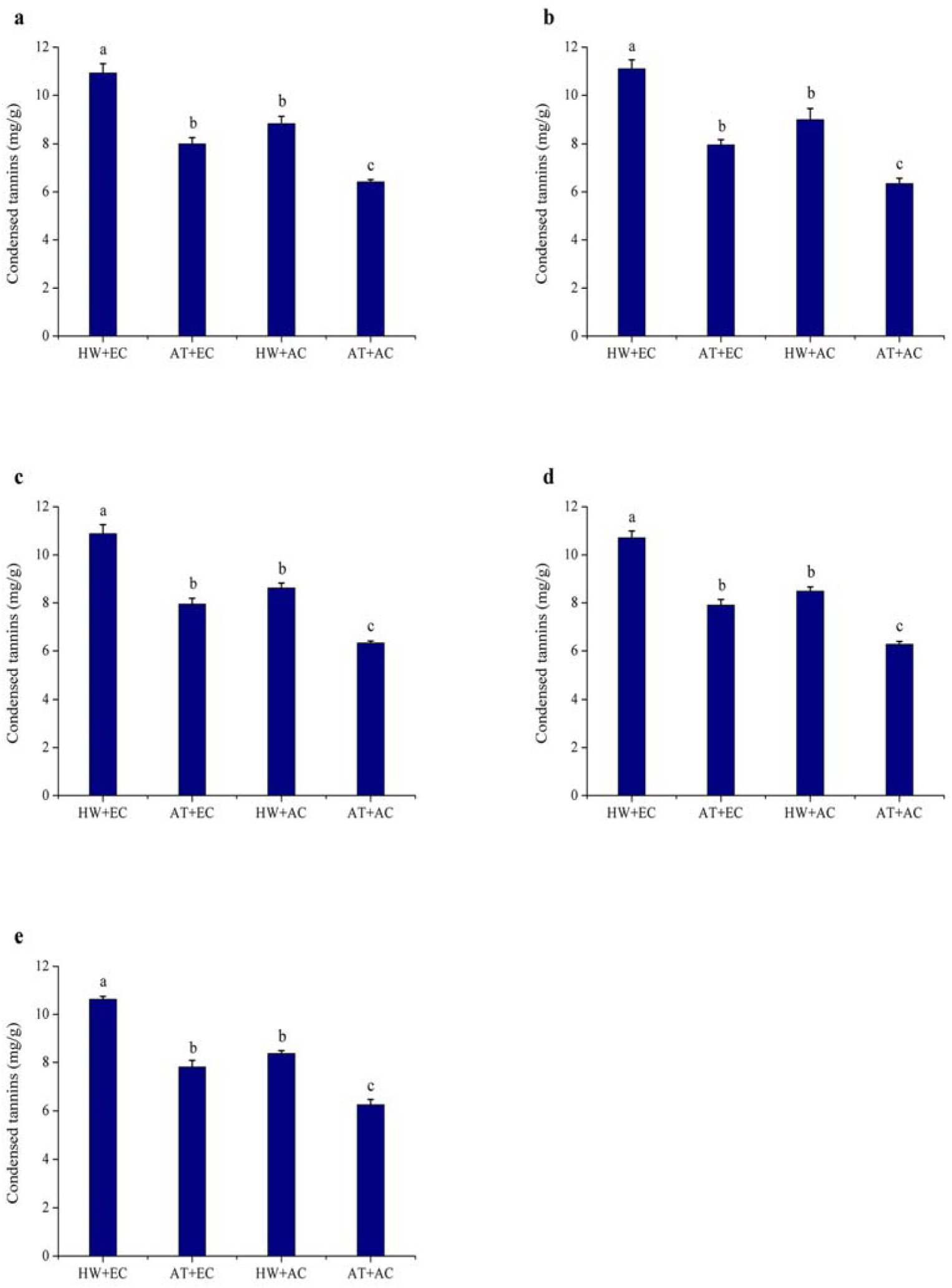
Condensed tannins concentration (±SE) of *Ambrosia artemisiifolia* under different stress conditions. Data represented by columns bearing the same letters were not significantly different (LSD, *P* = 0.05). A, B, C, D, and E represent the 1^st^, 2^nd^, 3^rd^, 4^th^, and 5^th^ measurements, respectively. AT denotes the normal temperature condition; HW denotes the heat wave condition. EC and AC represent elevated atmosphere CO_2_ concentration and ambient atmosphere CO_2_ concentration, respectively.

### Effect on the development period, female ratio, adult longevity, and female fecundity of *O. communa* after feeding on common ragweed grown under different treatment conditions

The development period of each stage, female ratio, adult longevity, and female fecundity of *O. communa* are presented in Tables 1 and 2. The first larval stage, the whole larval stage, the pupal stage, and the entire immature stage of *O. communa* were significantly increased when beetles fed on common ragweed grown under the high CO_2_concentration treatment. Furthermore, among the four treatments, *O. communa* that fed on common ragweed treated with a combination of high CO_2_concentration and heat wave (EC+HW) had the longest developmental period (*P*<0.05). There was no significant difference in terms of the female ratio among *O. communa* populations fed on common ragweed grown under the four treatments. However, compared with the AC+ AT treatment, the longevity of *O. communa* adults and the fecundity of females that fed on common ragweed treated with high CO_2_concentration were significantly decreased (*P*< 0.05). The longevity of the *O. communa* adults that fed on common ragweed treated under EC+HW conditions was the shortest, and the difference in female longevity between the EC+AT and EC+HW treatments was significant. For fecundity, irrespective of the treatment of heat wave alone (AC+HW), high concentration CO_2_ alone (EC+AT), and a combination of high concentration CO_2_ and heat waves (EC+HW), the number of eggs laid per female was significantly decreased compared with the control (AC+AT) treatment (*P* < 0.05).

**Table 1.**
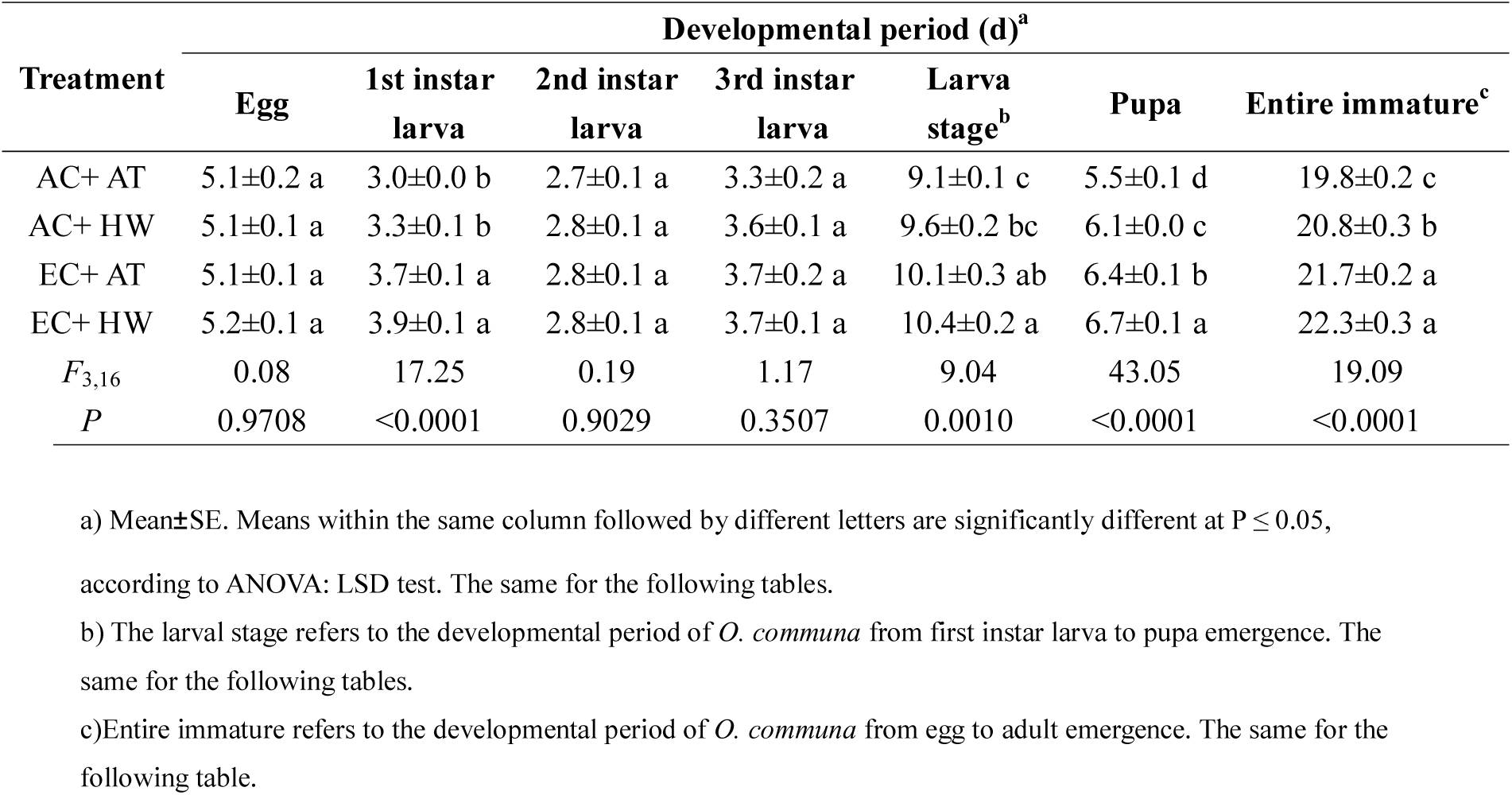
Effect of different stages of *Ophraella communa* by feeding on common ragweed after treatment on development period

| Treatment | Developmental period (d)<sup>a</sup> |  |  |  |  |  |  |
| --- | --- | --- | --- | --- | --- | --- | --- |
|  | Egg | 1st instar larva | 2nd instar larva | 3rd instar larva | Larva stage<sup>b</sup> | Pupa | Entire immature<sup>c</sup> |
| AC+ AT | 5.1±0.2 a | 3.0±0.0 b | 2.7±0.1 a | 3.3±0.2 a | 9.1±0.1 c | 5.5±0.1 d | 19.8±0.2 c |
| AC+ HW | 5.1±0.1 a | 3.3±0.1 b | 2.8±0.1 a | 3.6±0.1 a | 9.6±0.2 bc | 6.1±0.0 c | 20.8±0.3 b |
| EC+ AT | 5.1±0.1 a | 3.7±0.1 a | 2.8±0.1 a | 3.7±0.2 a | 10.1±0.3 ab | 6.4±0.1 b | 21.7±0.2 a |
| EC+ HW | 5.2±0.1 a | 3.9±0.1 a | 2.8±0.1 a | 3.7±0.1 a | 10.4±0.2 a | 6.7±0.1 a | 22.3±0.3 a |
| *F*<sub>3,16</sub> | 0.08 | 17.25 | 0.19 | 1.17 | 9.04 | 43.05 | 19.09 |
| *P* | 0.9708 | <0.0001 | 0.9029 | 0.3507 | 0.0010 | <0.0001 | <0.0001 |

a) Mean±SE. Means within the same column followed by different letters are significantly different at $P \leq 0.05$ ,
according to ANOVA: LSD test. The same for the following tables.
b) The larval stage refers to the developmental period of *O. communa* from first instar larva to pupa emergence. The same for the following tables.
c) Entire immature refers to the developmental period of *O. communa* from egg to adult emergence. The same for the following table.

**Table 2.** Effect on female ratio, longevity of adult, and fecundity of female *O. communa* after feeding on common ragweed grown under different treatment conditions.

| Treatment | Female ratio (%) | Female longevity (d) | Male longevity (d) | No. eggs laid per female |
| --- | --- | --- | --- | --- |
| AC+ AT | 53.7±0.7 a | 43.1±0.6 a | 56.6±1.0 a | 1137.0±12.8 a |
| AC+ HW | 51.8±3.1 a | 42.6±0.6 a | 58.6±0.3 a | 1070.6±7.6 b |
| EC+ AT | 50.8±1.8 a | 38.2±0.7 b | 53.7±1.1 b | 637.7±33.4 c |
| EC+ HW | 52.0±2.2 a | 32.7±1.2 c | 52.5±0.9 b | 581.0±23.9 c |
| *F*<sub>3,16</sub> | 0.31 | 34.90 | 9.87 | 173.41 |
| *P* | 0.8152 | <0.0001 | 0.0006 | <0.0001 |

a) Mean± SE. Means within the same column followed by different letters are significantly different at $P \leq 0.05$ as determined by an ANOVA: LSD test.
b) The larval stage refers to the developmental period of *O. communa* from first instar larva to pupa emergence.
c) Entire immature refers to the developmental period of *O. communa* from egg to adult emergence.

### Effect on the survival rate of different stages of *O. communa* after feeding on common ragweed grown under different treatments

As shown in Table 3, there was no significant difference in the survival rate of larvae when *O. communa* larvae were fed on common ragweed treated under the four stress conditions. However, the survival rate of pupa and the entire immature stage was significantly lower in *O. communa* fed on common ragweed treated under EC+HW than that of *O. communa* fed on common ragweed treated under AC+AT conditions.

**Table 3.** Effect on survival of *Ophraella communa* after feeding on common ragweed grown under different treatment conditions.

| Treatment | Survival rate (%) |  |  |  |  |  |  |
| --- | --- | --- | --- | --- | --- | --- | --- |
|  | Egg | 1<sup>st</sup>instarlarva | 2<sup>nd</sup>instarlarva | 3<sup>rd</sup>instarlarva | Larva stage | Pupa | Entire immature |
| AC+AT | 85.8±2.2a | 75.3±5.7a | 92.6±1.6a | 92.9±3.5a | 64.1±2.9a | 97.1±0.4a | 53.3±2.3a |
| AC+HW | 84.4±1.6a | 73.6±2.9a | 92.1±1.6a | 92.0±3.1a | 61.2±3.8a | 94.4±1.6ab | 48.5±2.8ab |
| EC+AT | 82.4±3.3a | 81.7±2.9a | 90.0±2.1a | 90.2±2.3a | 66.5±3.9a | 87.6±3.8b | 46.8±2.1ab |
| EC+HW | 81.5±2.3a | 77.4±5.8a | 85.6±5.4a | 88.2±4.7a | 64.8±5.1a | 88.5±3.7b | 43.7±3.4b |
| $F_{3,16}$ | 0.62 | 0.59 | 1.06 | 0.36 | 0.30 | 2.79 | 2.19 |
| $P$ | 0.6136 | 0.6287 | 0.3927 | 0.7850 | 0.8226 | 0.0745 | 0.1291 |

a) Mean±SE. Means within the same column followed by different letters are significantly different at $P \leq 0.05$ ,
according to ANOVA: LSD test. The same for the following tables.
b) The larval stage refers to the developmental period of *O. communa* from first instar larva to pupa emergence. The same for the following tables.
c) Entire immature refers to the developmental period of *O. communa* from egg to adult emergence.

### Effect on the life table parameters of *O. communa* after feeding on common ragweed grown under different treatment conditions

Compared with the control group, the intrinsic rate of increase (*r*) and net reproduction rate (*R*o) decreased significantly and the generation time (T) increased significantly in *O. commun*a fed on common ragweed grown under the other three treatments. There was no significant difference among *O. communa* fed on common ragweed grown under the four treatments. In addition, *O. communa* fed on common ragweed treated under EC+HW conditions had the lowest values of *r*, *Ro*, and, and had the longest generation time, which were 0.1340, 129.4, 1.1435, and 36.2, respectively (Table 4).

**Table 4.** Effect on the life table parameters of *Ophraella communa* after feeding on common ragweed grown under different treatment conditions.

| Treatment | Intrinsic rate of increase ($r$) | Net reproduction rate ($R_0$) | Generation time (T) | Finite rate of increase ($\lambda$) |
| --- | --- | --- | --- | --- |
| AC+ AT | 0.1821±0.0022 a | 324.6±19.6 a | 31.7±0.4 c | 1.3556±0.1554 a |
| AC+ HW | 0.1698±0.0018 b | 265.4±3.1 b | 32.9±0.3 b | 1.1851±0.0021 a |
| EC+ AT | 0.1511±0.0019 c | 159.9±8.6 c | 33.6±0.4 b | 1.1631±0.0022 a |
| EC+ HW | 0.1340±0.0026 d | 129.4±9.5 c | 36.2±0.3 a | 1.1435±0.0030 a |
| $F_{3,16}$ | 98.48 | 59.09 | 28.00 | 1.57 |
| $P$ | <0.0001 | <0.0001 | <0.0001 | 0.2356 |

a) Mean±SE. Means within the same column followed by different letters are significantly different at $P \leq 0.05$ , as
determined by an ANOVA: LSD test.
b) The larval stage refers to the developmental period of *O. communa* from first instar larva to pupa emergence.
c) Entire immature refers to the developmental period of *O. communa* from egg to adult emergence.

The maximum daily fecundity was 22.5, 23.4, 18.8, and 16.4 under AC+AT, AC+HW, EC+AT, and EC+HW, respectively; the peak fecundity occurred on days 29, 31, 32, and 42, respectively. The duration of the peak fecundity in the AC+AT treatment was longer and decreased gradually, while that of AC+HW and EC+AT treatments decreased rapidly after reaching the peak reproductive stage. Compared with the other three treatment groups, the peak reproductive stage of the EC+HW treatment group appeared later, decreased rapidly after reaching the peak, rose again, and then decreased gradually (Fig. 1). The age-specific survivorship of each treatment decreased rapidly in 25 days, with the survivorship for AC+AT, AC+ HW, EC+ AT, and EC+ HW decreasing rapidly after 39, 34, 34, and 34 days, respectively (Fig. 3).

**Fig.3.**
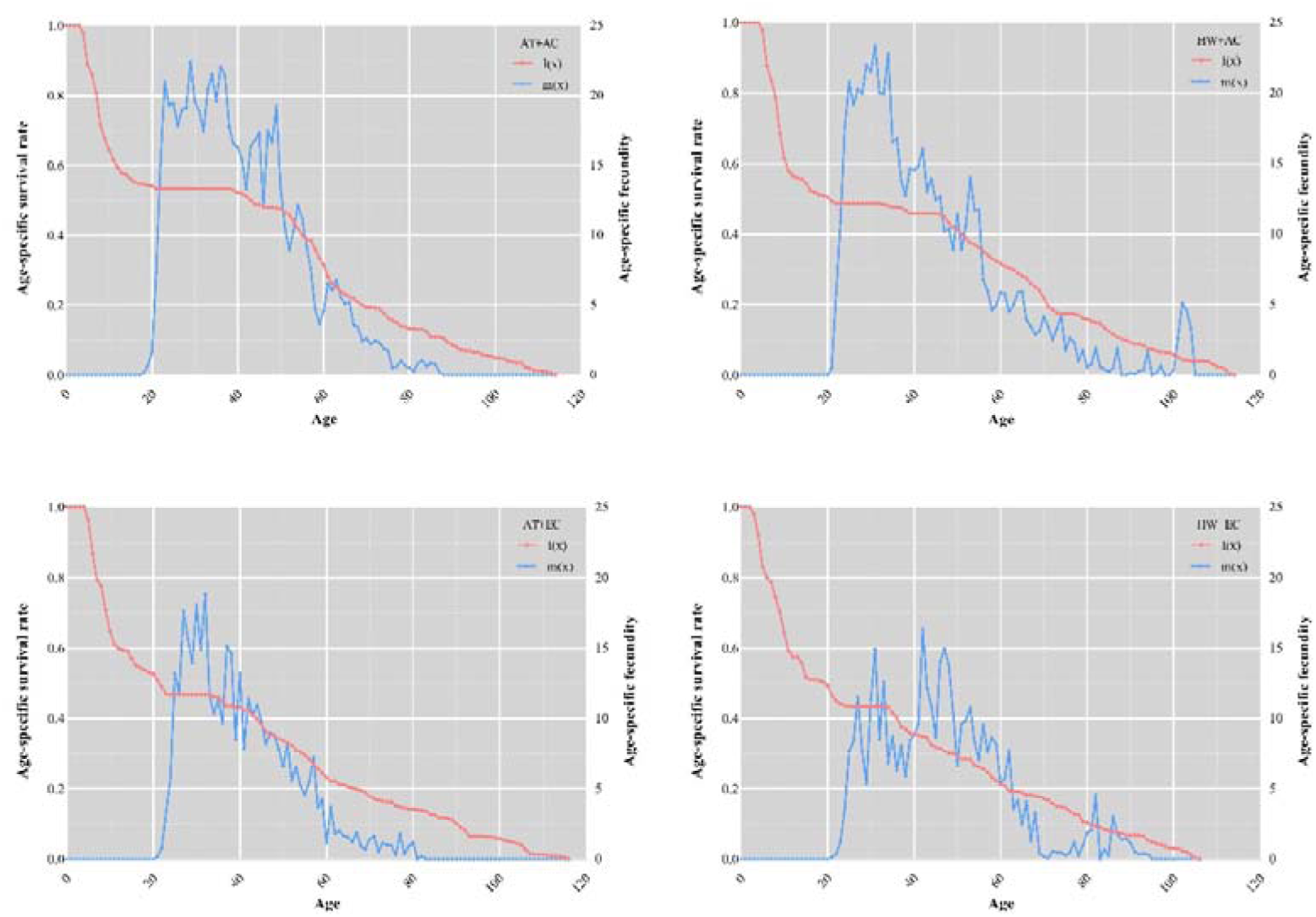
Effect on age-specific survivorship and fecundity of *Ophraella communa* after feeding on common ragweed grown under different treatment conditions. AT denotes the normal temperature condition; HW denotes the heat wave condition. EC and AC represent elevated atmosphere CO_2_ concentration and ambient atmosphere CO_2_ concentration, respectively.

### Effect on enzyme activity of *O. communa* after feeding on common ragweed grown under different treatment conditions

Compared with common ragweed grown under AC+AT conditions, the enzyme activities (including four detoxification enzymes and two ingestion enzymes) of the adults were changed after feeding on ragweed grown under EC+HW, EC+AT, and AC+HW conditions; the enzyme activities of catalase (CAT), superoxide dismutase (SOD), acetylcholinesterase (AChE), carboxylesterase (CarE), tryphin, and lipase increased. In all experimental conditions, the enzyme activity of female adults was higher than that of males, and the different experimental conditions had no significant effect on this result. A comparison of the enzyme activities under the four experimental conditions found that the changes in enzyme activities in the adults fed on common ragweed grown under EC+ HW conditions were the most significant, having the greatest impact on enzyme activities (Fig. 4).

**Fig.4.**
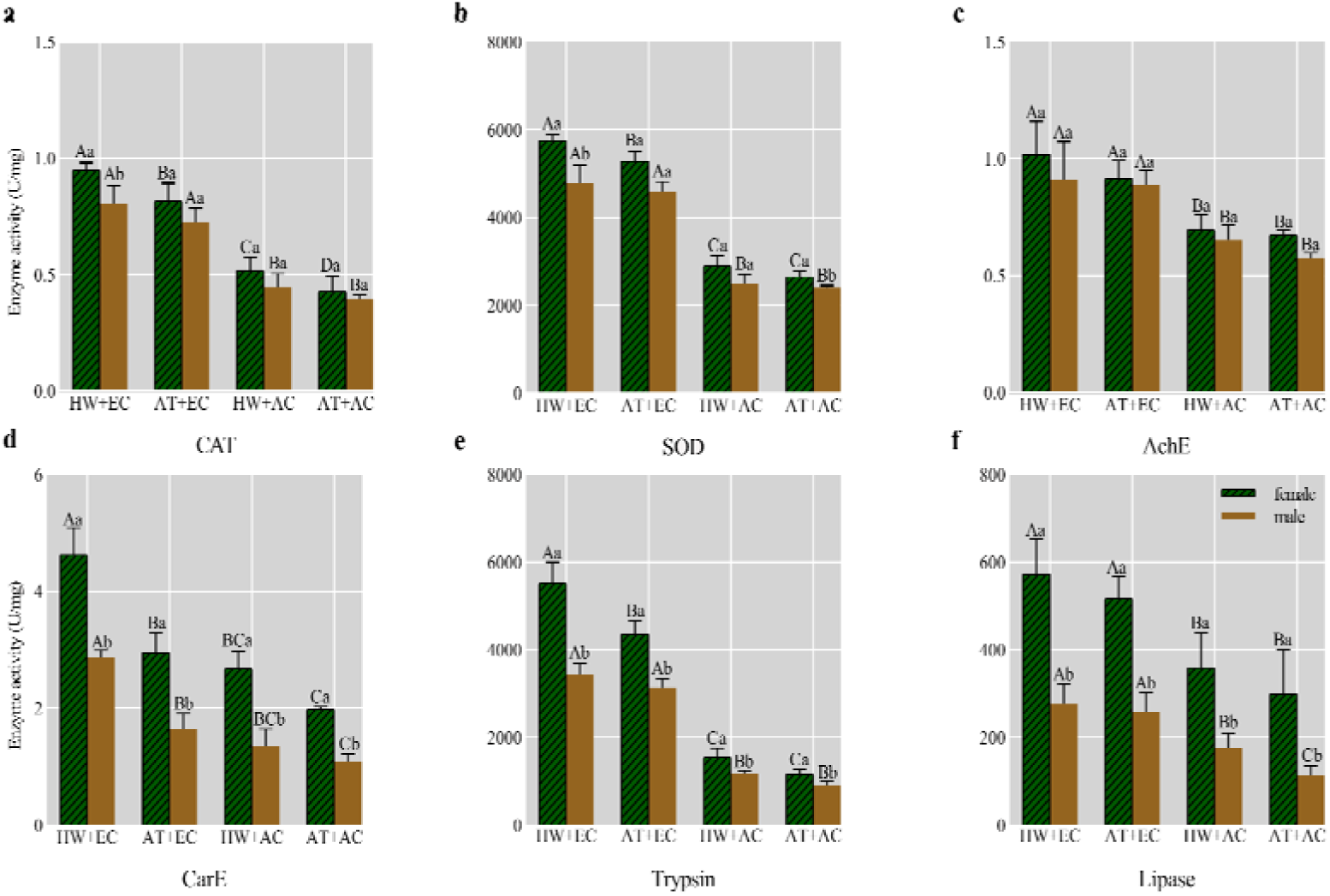
Effect on seven enzyme activities of *Ophraella communa* after feeding on common ragweed grown under different treatment conditions. AT denotes the normal temperature condition; HW denotes the heat wave condition. EC and AC represent elevated atmosphere CO_2_ concentration and ambient atmosphere CO_2_ concentration, respectively.

### Metabonomic profile of common ragweed after high CO_2_ concentration and heat wave treatment

A total of 589 metabolites were measured from the leaves of plants under AC+AT and EC+HW conditions. Compared with the common ragweed growing in the AC+AT treatment, there were 163 different metabolites in the leaves of the common ragweed under the EC+HW treatment, 88 of which were significantly up regulated (VIP ≥ 1, FC ≥ 2) and 75 of which were significantly down regulated (VIP ≥ 1, FC ≤ 0.5, Fig.5a). In addition, the content of N-(p-coumaroyl) serotonin in common ragweed leaves under the EC+HW treatment was 2264.4-fold higher than that in the AC+AT treatment.

**Fig.5.**
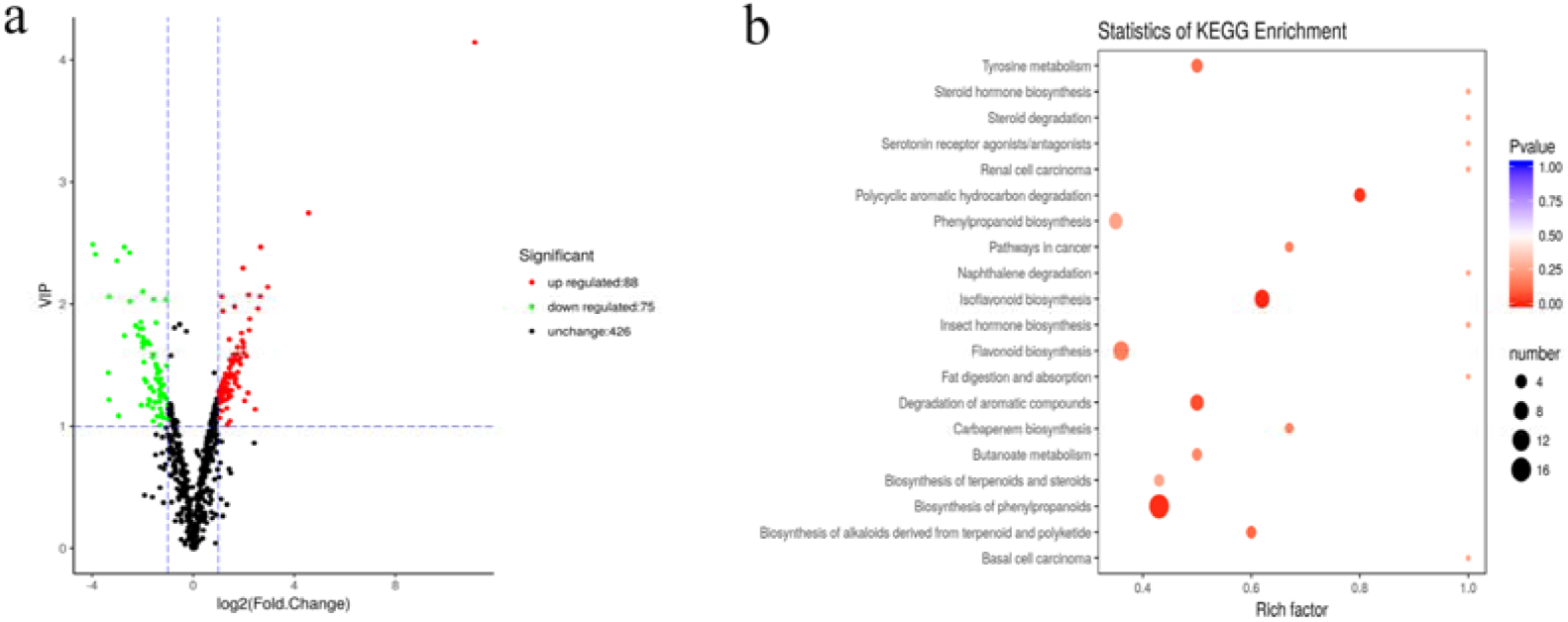
a: Volcano plot of differentially expressed metabolites; b: Statistics of KEGG enrichment. Leaves of plants under AC+AT and EC+HW conditions.

The classification results of KEGG annotation of different metabolites are shown in Supplementary Fig. 4. There were163 different metabolites annotated in 116 different pathway species, including 7 major pathway categories, most of which were concentrated in metabolism. Among them, 36 metabolites belonged to the metabolic pathway and 27 belonged to the biosynthesis of secondary metabolites, accounting for 54.55% and 40.91%, respectively. After enrichment of the KEGG pathway, it was found that the pathways of differential metabolites were mainly concentrated inflavonoid biosynthesis, isoflavonoid biosynthesis, and phenylpropanoid biosynthesis (Fig.5b).

#### Bioassay test

Eight significantly up regulated substances (Cryptochlorogenic acid, N-(p-Cinnamoyl) serotonin, Glycitin, Phe-Phe, Fumaric acid, A-Ketoglutaric acid, Calycosin, and Glycitein) were used to perform the bioassay test. Compared with the control group, feeding on common ragweed leaves treated with cryptochlorogenic acid (2000 ppm,1000 ppm, and 500 ppm), N-(p-Cinnamoyl) serotonin (2000 ppm, 1500 ppm, and 1000 ppm), Glycitin (1500 ppm and 1000 ppm), Phe-Phe (1000 ppm and 500 ppm), Fumaric acid (500 ppm), or A-Ketoglutaric acid (500 ppm) significantly increased the mortality of the leaf beetles. In addition, the leaf beetles feeding on Calycosin- and Glycitein-treated leaves had higher mortality than the control group, although there was no significant difference (Table 5). For Glycitin, Phe-Phe, Fumaric acid, and A-Ketoglutaric acid, the trend of mortality showed a phenomenon of optimal lethal concentration, i.e., with the increase of the concentration of the substance provided, the mortality of the beetle increased gradually, but when higher concentration of substance was continually provided, mortality began to decrease.

**Table 5.**
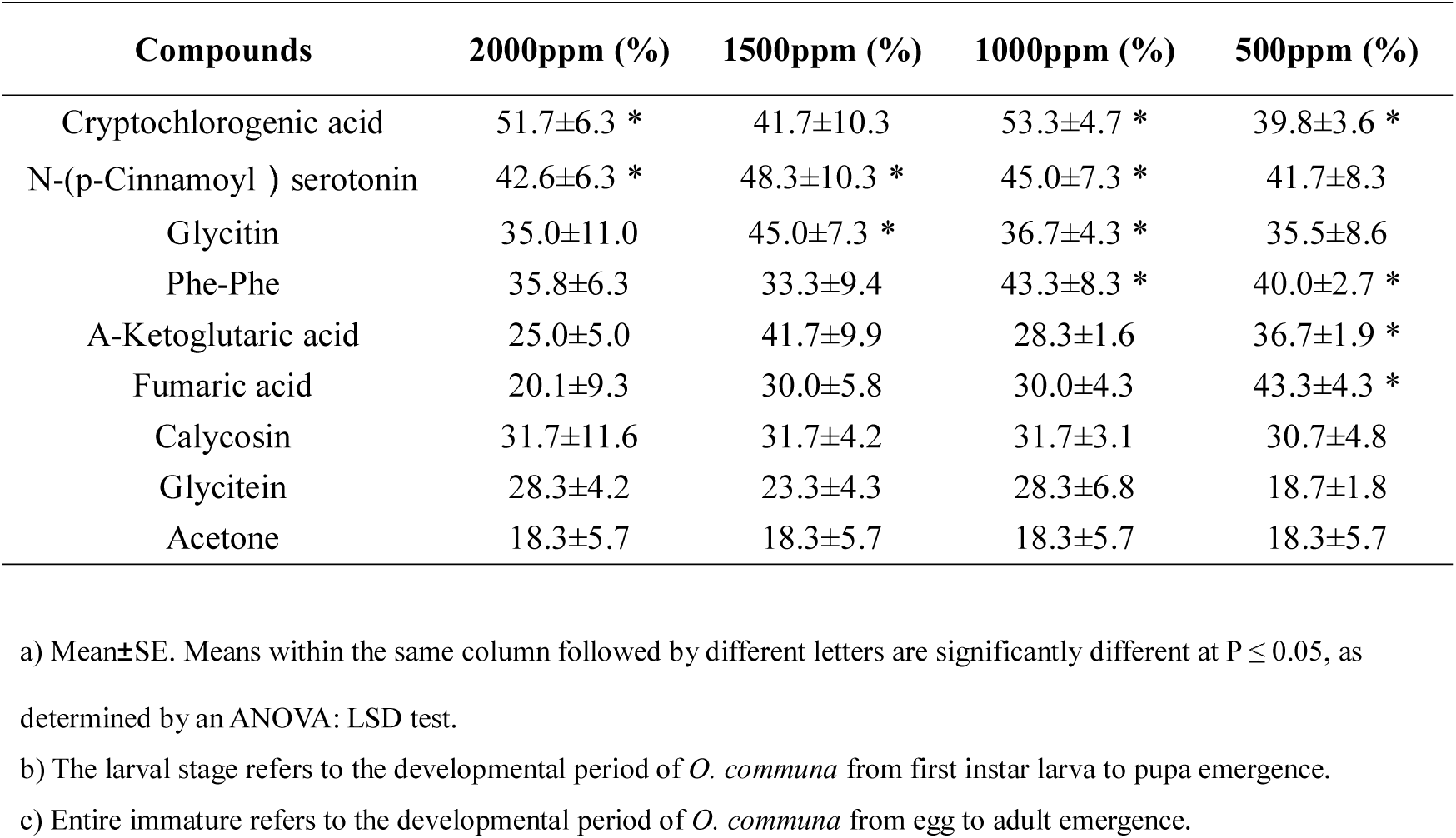
Effect on immature death rate of *Ophraella communa* after feeding on common ragweed grown under different treatment conditions.

| Compounds | 2000ppm (%) | 1500ppm (%) | 1000ppm (%) | 500ppm (%) |
| --- | --- | --- | --- | --- |
| Cryptochlorogenic acid | 51.7±6.3 * | 41.7±10.3 | 53.3±4.7 * | 39.8±3.6 * |
| N-(p-Cinnamoyl ) serotonin | 42.6±6.3 * | 48.3±10.3 * | 45.0±7.3 * | 41.7±8.3 |
| Glycitin | 35.0±11.0 | 45.0±7.3 * | 36.7±4.3 * | 35.5±8.6 |
| Phe-Phe | 35.8±6.3 | 33.3±9.4 | 43.3±8.3 * | 40.0±2.7 * |
| A-Ketoglutaric acid | 25.0±5.0 | 41.7±9.9 | 28.3±1.6 | 36.7±1.9 * |
| Fumaric acid | 20.1±9.3 | 30.0±5.8 | 30.0±4.3 | 43.3±4.3 * |
| Calycosin | 31.7±11.6 | 31.7±4.2 | 31.7±3.1 | 30.7±4.8 |
| Glycitein | 28.3±4.2 | 23.3±4.3 | 28.3±6.8 | 18.7±1.8 |
| Acetone | 18.3±5.7 | 18.3±5.7 | 18.3±5.7 | 18.3±5.7 |

a) Mean±SE. Means within the same column followed by different letters are significantly different at $P \leq 0.05$ , as
determined by an ANOVA: LSD test.
b) The larval stage refers to the developmental period of *O. communa* from first instar larva to pupa emergence.
c) Entire immature refers to the developmental period of *O. communa* from egg to adult emergence.

## Discussion

Climate change is the most important factor influencing natural ecosystems and organisms (Jeong et al., 2018; Howden et al., 2007).The elevatedCO_2_ concentration has attracted attention worldwide, and it is expected to drive wide spread increases in extreme heat events this century (Skinneret al., 2018).In this study, we focused on the effect of elevated CO_2_and heat waves on common ragweed secondary metabolism change and on the performance of the common ragweed-specialist herbivorous insect *O. communa* through metabolomics and a two-sex life table, respectively, subsequently demonstrating the interaction of common ragweed–leaf beetle under a climate change context and providing guidance or common ragweed biocontrol in the future.

Organisms are more sensitive to abrupt than to gradual change, and extreme climate events are considered to be the main instigators of evolution (Combes, 2008; Gutschick & BassiriRad, 2003).In addition, climate change will result in a series of chain reactions, affecting organisms at all trophic levels (Rouault et al., 2006).On the other hand, extreme climatic events can enhance invasion processes, from initial introduction through to establishment and spread (Diez et al., 2012; Walther et al., 2009). Compared with native plants, invasive plant species may show a high capacity to cope with environmental changes due to greater physiological, phenotypic, and ecological plasticity (Davidson et al., 2011; Van Kleunen et al., 2011). Kiesel (2014) also reported that rapid adaptation to changes in climate may in fact be key to the success of invasive plants. Recently, Sun et al. (2020) demonstrated, by DNA sequencing and phenotypic observation, that common ragweed populations can rapidly evolve within a single generation in response to climate change.

Studies have reported that elevated CO_2_hasa beneficial impact on many aspects ofC3 plants, such as growth rate, biomass, and photosynthetic rate (Obermeieret al., 2017; Specht et al., 2014; Rodgers et al., 1983; Wong, 1979). In addition, the carbon-nutrient balance theory and the growth-differentiation balance model predict that plants will prioritize photosynthesis and accumulation of carbohydrates over growth under elevated atmospheric CO_2_ concentration conditions, and that excess carbon will be allocated to the production of carbon-based plant secondary metabolites (Herms & Mattson, 1992; Bryant et al., 1983).Thus, changes in environmental conditions will affect the secondary metabolic pathway of plants, which can change the type and content of secondary metabolites and activate the plant’s defense response in various ways (Leonardo & Norberto 2007).For example, high concentration of CO_2_increasedthe tannin and gossypol content of *Bt* cotton (Chen et al., 2005;Coviella, 2002; Coviella et al., 2000) and the content of saponin in alfalfa (Agrell et al., 2004).Heavy metals significantly stimulated the accumulation of saponins, phenolic compounds, and flavonoids in *Robiniapseu doacacia* leaves and stems, while alkaloid compounds increased in leaves and decreased in stems (Zhao et al., 2016).Plant secondary metabolites play important roles in various aspects of plant biology, including regulation of growth (Mikkelsen et al., 2015) and the suppression (through allelochemicals) of competing plants (Inderjit, 1996).In addition, numerous studies have reported that plant secondary metabolites are essential for protecting plants from adverse environments(e.g., UV light heavy metals, and heat) (Zhao et al., 2016; Veteli et al., 2003).Previous studies have reported that high levels of CO_2_ induce plants to increase their accumulation of carbon-based secondary compounds, while gentle warming decreases the production of secondary compounds, counteracting the defensive effects of plants induced by high CO_2_ concentrations (Veteli et al., 2002; Lincoln et al., 1993).In the present study, however, elevated CO_2_alone,heat wave alone, and a combination of elevated CO_2_ and heat wave remarkably enhanced the concentrations of total phenolics and condensed tannins in common ragweed. In addition, unlike gentle warming, which counteracts the plant defensive response induced by high CO_2_, the heat wave significantly intensified the defensive response of plants, resulting in more phenolics and condensed tannins. This suggests that there is a significant synergetic relationship between elevated atmospheric CO_2_ concentration and heat waves, which stimulate more intense defensive responses in plants; increased secondary compound accumulation in plants is likely to happen under future climate change conditions. The metabolomic profile results showed that, compared with the control treatment, the concentrations of 163 metabolites changed significantly in the leaves of common ragweed after combined treatment with elevated CO_2_ and heat waves (Fig.5). Moreover, the content of N-(p-coumaroyl) serotonin in common ragweed leaves under the EC+HW treatment was2264.4-foldhigher than that in the AC+AT treatment. This large-scale change in secondary metabolites demonstrated that common ragweed had a strong ability to activate its defense system and adjust the secondary metabolite pathways to resist the unfavorable environment, which will facilitate common ragweed to better adapt to future climate change and develop its invasion potential.

Another important function of plant secondary metabolites is defense against herbivores (Jiaet al., 2014; Rausher, 2001). Previous studies have reported that secondary metabolites like alkaloids, flavonol glycosides, gossypol, and tannins are toxic to insects, resulting in slow growth of insects, inhibition of molting, and decreased fecundity (Stipanovic et al., 2006; Canals et al., 2005; Lindroth et al., 1997; Traw et al., 1996).

Thus, climate change likely would also influence insect performance and affect plant-herbivore interactions through changes in the levels of secondary metabolites in plants. In the present study, elevated CO_2_ alone, heat wave alone, and a combination of elevated CO_2_ and heat wave increased *O. communa* developmental period (larva and pupa stages) but decreased their survival rate, adult longevity, and fecundity. Compared with the control treatment, the intrinsic rate of increase and net reproduction rates ignificantly increased but generation time significantly decreased in the other three treatments. This signifies that feeding on common ragweed grown under elevated CO_2_ and heat wave conditions had a profound impact on the development and reproduction of *O. communa* populations. Detoxifying and digestive enzymes are often used as important criteria for measuring physiological changes in insects (Li et al., 2017; Zhu-Salzman &Zeng, 2015). Enzyme activity results showed that, irrespective of sex, the activities of six important enzymes in beetles fed on common ragweed treated with elevated CO_2_were significantly upregulated. In addition, beetles fed on common ragweed treated with elevated CO_2_and heat wave had high EST enzyme activities. This is because elevated CO_2_ and heat wave increased the concentration of secondary metabolites in common ragweed (e.g., flavonoid and tannins; Fig. 1, Fig. 2, Fig. 5). Detoxification and digestion require the consumption of many nutrients and energy in insects (Schoonhoven & Meerman, 1978). Therefore, when *O. communa* responds to the defensive reaction of common ragweed induced by high concentrations of CO_2_ and heat waves, it needs to allocate more energy for detoxification and digestion, which will prolong the development time and reduce spawning. The above results—including the development period, fecundity, survival rate, life table parameter, and enzyme activity of *O. communa—*show that heat waves strengthen the defense effect of common ragweed against herbivores through changes in the levels of secondary metabolites under elevatedCO_2_conditions. Based on the results of the metabolomics, we selected eight significantly upregulated substances for a bioassay test. The results showed that the mortality rates of *O. communa* (immature stage) under six substances were significantly higher than those in the control group, demonstrating that the combination of elevated CO_2_ and heat wave-induced secondary metabolites was unfavorable for the development of *O. communa*. Consistent with our results, Jamieson et al. (2015) reported that warming induced higher levels of condensed tannin and phenolic glycoside in *Populus tremuloides* and markedly increased larval development times and decreased larval weight and food conversion efficiency. Thus, climate change induced changes in phytochemicals leading to corresponding host species-specific changes in insect performance (Jamieson et al., 2015). On the other hand, under field conditions, changes in the secondary metabolite content of common ragweed prolonged the development time, meaning that the exposure time of *O. communa* to natural enemies will be increased and the population size of *O. communa* will be decreased. This will bring challenges to the biological control of common ragweed in the future.

As a worldwide allergenic, invasive weed, common ragweed has brought great trouble to agriculture and human health. Therefore, prediction of the performance of common ragweed under a climatic change trend is beneficial to prevent its distribution and reduce its harm to human health. It is indisputable that the concentration of CO_2_ in the atmosphere will rise in the future and will increase the global temperature and drive the frequency and intensity of heat waves (Skinner et al., 2018). Previous studies have demonstrated that elevated CO_2_ and warming will boost common ragweed biomass, reproduction, pollen production, and allergenic ability (Sun et al., 2020; Rogers et al., 2006; Stinson & Bazzaz, 2006; Stinson et al.,2006). Sun et al. (2020) also demonstrated that common ragweed populations can rapidly evolve in response to climate change within a single generation, which will aggravate the impact of its distribution and damage in the future. Moreover, our results demonstrated that elevated CO_2_ and more extreme heat events can help common ragweed improve photosynthesis, increase total leaf area, and increase growth (Supplementary Fig.1,2, and 3), which will undoubtedly increase the competitive invasion ability of this species in the future. On the other hand, elevated CO_2_ and heat wave-induced accumulation of secondary metabolites in common ragweed had an unfavorable impact on the development of *O. communa*. Under elevated CO_2_ conditions, heat waves can promote common ragweed’s ability to adapt to climate change and defense against herbivores, making it more difficult to prevent and control common ragweed in the future. On the other hand, we can also conclude that heat wave event will relieve harm of *A. artemisiifolia* to human under elevated CO_2_ in the future.

## Materials and methods

### Part 1: Effects of high CO_2_concentration and heat wave on the performance of *A. artemisiifolia* and *O. communa*

#### Control of CO_2_ concentrations

Experiments were conducted in four climate-controlled chambers (PRX-450C-CO_2_; Ningbo Haishu Saifu Experimental Instrument Factory, Ningbo, China) located at the Langfang Experimental Farm of Hebei Province, China ’E 116°36’46“, N39°30’36“, at an altitude of 27 m). Growth chambers were used rather than open-top chambers or Free-Air CO_2_ Enrichment (FACE), to provide stable CO_2_, light, and humidity for continuous treatment effect throughout the experimental period. The CO_2_ concentrations were set at the current atmospheric CO_2_ concentration (370 ppm) and at an elevated CO_2_ concentration (700 ppm, which is predicted for the end of this century) (IPCC, 2013). Two blocks were used for each of the two CO_2_ treatments, and each block contained one pair of climate-controlled chambers (one with ambient CO_2_ and one with elevated CO_2_) (Sun et al. 2018). The CO_2_ concentration in each chamber was maintained by continual flushing with CO_2_-free air and then re-injecting CO_2_ from the cylinders to maintain the appropriate CO_2_ concentration. CO_2_ concentrations in all chambers were monitored using an absolute infrared gas analyzer (MSA Instruments, Pittsburgh, PA, USA).

#### Host plants

*Ambrosia artemisiifolia* seeds used in this study were collected from the Langfang Experimental Farm in the last third of October in 2014. The seeds were then stored in a freezer at 4°C. Adequately stored seeds were germinated in a greenhouse in late April 2015. When the seedlings reached a height of approximately 10 cm, seedlings were thinned to one per pot in all treatments and were transplanted into bigger pots (21 cm diameter and 17 cm height) containing soil (Chen et al., 2018). The pots were filled with vermiculite, nutrient soil, and natural soil in the volumetric ratio of 1:2:3. As experiments only examined the seedlings over 10 weeks, the plastic pot size was not a limiting factor. Each pot was fertilized weekly with a commercial fertilizer (Stanley Agriculture Group Co., Ltd, Linyi, China) and irrigated with 1 L of water every 3days. For the experiment, twenty potted plants were evenly spaced in two climate-controlled chambers, one with 370 ppm and the other with 700ppm CO_2_ concentration, respectively. Apart from the differing CO_2_ concentrations, the growth temperature conditions for the *A. artemisiifolia* seedlings before heat wave stresses were as follows: day/night temperatures were maintained at 30/25°C for 14/10 h, alongside a photoperiod of 14:10 L:D; a 70±5% relative humidity (RH)and a mean photosynthetic photon flux density (PPFD) of 600 μmol photons m^−2^ s^−1^ were maintained at all time. To avoid position effects, pots were rotated randomly inside each chamber and moved from one chamber to the other every 5 days, maintaining the corresponding CO_2_ treatment throughout the experiment.

#### Heat wave treatment

At 70 days after sowing, ragweed seedlings were subjected to a heat wave stress treatment. As the duration of heat wave events are more than five consecutive days (García-Herrera et al., 2010), the time of heat wave treatment was 5 days in this study. Half of each group of seedlings in the two CO_2_ concentration chambers described above were selected randomly for further cultivation under the original conditions, while the other half were transferred to two other growth chambers (one with 370 ppm and the other with 700 ppm of CO_2_) to undergo heat wave treatment. Therefore, there were 10 potted seedlings in each of the four growth chambers, which were provided the following treatment conditions: (1) ambient atmosphere CO_2_ concentration and ambient temperature (370 ppm, 30/25°C for 14/10 h in one day), acting as the control (AC+ AT); (2) ambient atmosphere CO_2_ concentration and heat wave (370 ppm, 40°C for 8 h, and 30/25°Cfor 6/10 h in one day), acting as the heat wave treatment (AC+HW); (3) elevated atmospheric CO_2_ concentration and ambient temperature (700 ppm, 30/25°Cfor 14/10 h in one day), acting as the elevated CO_2_ treatment (EC+AT); (4) elevated atmospheric CO_2_ concentration and heat wave (700 ppm, 40°C for 8 h, and 30/25°Cfor 6/10 h in one day), acting as the combination treatment of elevated CO_2_ and heat wave (EC+HW). The heat wave stress was imposed by increasing the chamber temperature to 40°C during the daytime for 8 h daily for 5 days. Each day, plants received 8 h of heat wave, starting at 9:00 and ending at 17:00, and the rest of the 24 h period remained at the same temperature as that of the control treatment (i.e., day temperatures from 6:00 to 9:00 and 17:00 to 20:00 were 30°C, and night temperatures from 20:00 to 5:00 were 25°C). The plants were watered and fertilized the same as before. After the 5-day heat wave treatment, heat wave-treatment chambers were returned to the same air temperature as that of the ambient treatment (30/25°Cfor 14/10 h) (Tripathee, 2008).

#### Growth data collection

Plant height and CO_2_ assimilation were measured five times: at the end of day 0, 5, 10, 15, and 20 post-heat wave treatment. The total leaf area of plants was measured on day 20 after heat wave treatment. CO_2_ assimilation was measured on single, young, fully expanded leaves using a portable photosynthesis system (HCM-1000; Walz, Germany) equipped with a 6cm^2^ leaf chamber. Measurements were made at ambient light intensity within 1 min of insertion of leaves into the environmentally controlled cuvette and after stabilization of CO_2_ flux, to ensure that photosynthetic response reflected conditions within the growth chambers. Growth rates were measured at a leaf C for the ambient air temperature treatment and at 40° between 09:00 and 12:00, at the same CO_2_ levels as that of the treatment conditions (370 or 700 ppm). To prevent desiccation of leaves during measurement, the air stream entering the chamber was humidified, and the measurements were made rapidly. Total leaf area was determined using a portable leaf area meter (model CI-202, CID, Inc., Camas, WA, USA). Plant height was measured with a tape measure from the apex of the plant to the soil surface. Five samples were randomly selected for measurement per treatment.

#### Plant material sampling and foliar chemistry

Leaves from plants in the treatments were sampled at five times over the course of the experiment, and five replicates were taken at each sampling event. Samples were collected at the same time as that when the plant measurements described above were undertaken. Sampled leaves were randomly selected and, after being picked, were immediately frozen in liquid nitrogen and stored at -80° Before chemical analysis of leaf tissue, leaves were dried in a circulating air oven at 68° ground in liquid nitrogen, and kept in plastic containers that were placed in a desiccator and stored in a C until time of use.

Extraction and quantification of total phenolic concentration followed the methodology of Ibrahim and Jaafar (2012). Briefly, finely ground leaf tissues (0.25 mm fragments) of standard weight (0.1 g) were extracted with 80% ethanol (10 mL) in an orbital shaker for 120 minutes at 50° mixture was then filtered (Whatman™ No. 1), and the filtrate was used for the quantification of total phenolics. Folin-Ciocalteu reagent (diluted 10-fold) was used to determine the total phenolic content of the leaf samples. Sample extract (200 µL, after filtration) was mixed with Folin-Ciocalteu reagent (1.5 mL) and allowed to stand at 22°C for 5 min before adding NaNO_3_ solution (1.5 mL, 60 g L^−1^). After 2h at 22°C, the absorbance at 725 nm was measured. The results were expressed as mg g^−1^ gallic acid equivalent (mg GAE g^−1^ dry sample).

Concentrations of condensed tannin were measured using the acid butanol method of Kanerva and Smolander (2008). Approximately, 0.1 g of ground sample was extracted with 1 mL of ice-cold 70% acetone containing 10 mM ascorbic acid at 4°C for 30 min. After centrifugation, the supernatants were pooled. After diluting the pooled sample supernatants with 350 µL of ascorbic acid, a 150-µL aliquot was reacted with 3.0 mL of 19:1 (v/v) N-butanol: concentrated HCl and 100 µL of 0.02 g mL^-1^ FeNH_4_(SO_4_)2·12H_2_O in 2 N HCl and incubated in a water bath (at 100°C) for 50 min. The absorbance of the solution at 550 nm was measured to quantify the condensed tannins. The reported total phenolics and condensed tannin concentrations are the means of five replicates.

#### Insects

More than 1,000 *O. communa* adults were collected from the town of Dajing (28°56′26″N, 113°14′38″E) in Miluo County, Yueyang City, Hunan Province, China, on June 24, 2014. Colonies of the beetle were maintained on *A. artemisiifolia* plants in the laboratory at 27±1°C at the Langfang Experimental Farm of Hebei Province, China. Pupae were collected and stored in a transparent plastic box (19 × 12 × 6 cm) and kept in an air-conditioned laboratory at 28±1°C. Newly emerged adults were collected, and males and females were separately held on potted *A. artemisiifolia* plants in cages (40 × 40 × 60 cm) in the same laboratory, at a density of 20 adults per plant and one plant per cage. Two-day-old adults were used for the experiments.

#### Development and survivorship of immature *O. communa*

The development and survivorship of immature *O. communa* were measured following the methodology of Zhou et al. (2010). *O. communa* adults from the above culture were paired, and each pair was placed on apotted fresh common ragweed plant in cages (60 × 60 × 80 cm) for oviposition. The potted plants with<12-h-old *O. communa* eggs were placed in a plastic basin (50 × 30 cm) to absorb water. Plants that had at least 80 eggs were placed randomly into the four environmental chambers (PRY-450D; Ningbo Haishu Aifu Experimental Equipment, Zhejiang, China), representing the four treatments, as described above. Ten plants with eggs were used in each treatment, and all environmental chambers were set at an RH of70±5% and a photoperiod of 14:10 L:D. Eggs were checked daily until all had hatched. The neonates were kept on the same common ragweed plant until pupation. Pupae were detached from the leaves, placed in an unsealed cuvette individually, and checked daily until they emerged as adults. Each treatment was repeated five times in the same environmental chamber, and the next cohort of eggs was placed when all larvae in the previous treatment became pupae. Survival rates and developmental periods at different developmental stages were recorded. All emerged adults were paired, and if the number of males was greater than that of females, new females were introduced to record the lifespan of males; if the number of females was greater than that of males, new males were introduced to record the egg production and lifespan of females.

#### Longevity and fecundity of leaf beetle adults

Twigs of common ragweed were inserted into plastic bottles (3 × 5 cm, diameter × height) filled with water and with a 0.8-cm-diameter hole in the lid to hold the twig. Newly emerged *O. communa* adults in the four treatments were mated, and each pair was placed on fresh common ragweed twigs. Twigs with beetles were placed in transparent plastic boxes (19 × 12 × 6 cm) covered with a mesh to prevent beetles from escaping. These plastic boxes with *O. communa* adults and *A. artemisiifolia* twigs were placed in environmental chambers under the same conditions as their immature stages. Fresh twigs were changed daily, and the eggs of *O. communa* were counted. The observation ended when the female adult died. If the male died earlier, another newly emerged male was added. Pre-oviposition and oviposition periods, longevity, and number of eggs laid per female were recorded.

#### Determination of enzyme activity

Catalase (CAT) activity was determined by the method based on Aebi (1984). The activity of superoxide dismutase (SOD) was determined using the method of Marklund and Marklund (1974). Acetylcholinesterase (AChE) activity was determined using acetylcholinesterase iodide (atchl) as a substrate (Ellman et al., 1961). Carboxylesterase (CarE) activity was determined using the methodology of Kandil et al. (2017). According to the method of Visweshwar et al. (2015), Nα - benzoyl-L-arg-*p*-nitroanilide (Ba*p*NA; Sigma Aldrich) was used as the specific substrate to determine the activity of trypsin.

### Part 2: Metabonomic profile of common ragweed after high CO_2_concentration and heat wave treatment

Based on the results of *A. artemisiifolia* foliar chemistry and *O. communa* life table, we collected common ragweed leaf samples at the end of the control treatment (AC+AT) and the combination treatment of elevated CO_2_ and heat wave (EC+HW) to analyze changes in all metabolites using metabolomics methods.

#### Leaf sampling and preparation

Leaves from plants in the treatments were sampled 5 days after the heat wave treatment. Sampled leaves were randomly selected and were immediately frozen in liquid nitrogen after being picked and stored at -80. Each treatment had three replicates.

The metabolomics methods were as described by Chen et al. (2013). The freeze-dried leaves were crushed (30Hz,1.5min) to powder by a mixer mill (MM 400; Retsch). Then, 100g of the leaf power was dissolved in methanol (Chromatographic purity, Merck) extract. The solution was placed in a refrigerator at 4°C overnight, and the extraction rate was improved by vortexing three times. After centrifugation (rotation speed 10,000 × *g*, 10 min), the supernatant was aspirated, and the sample was filtered with a microporous membrane (0.22 μm pore size) and stored in the injection bottle for liquid chromatography (LC)-mass spectrometry (MS)/MS analysis.

Additionally, to ensure data quality for metabolic profiling, quality control (QC) samples were prepared—by pooling aliquots of all samples that were representative of those under analysis—and used for data normalization. QC samples were prepared and analyzed using the same procedure as that for the experimental samples in each batch.

#### Ultra-performance LC-MS/MS analysis

The metabolomics methods used were as described by Chen et al. (2013). The LC-MS/MS portion of the platform was based on a UHPLC system (1290 series; Agilent Technologies) equipped with an ACQUITY UPLC BEH Amide column (1.7 μm, 2.1 mm 100 mm, Wasters) and a triple quadruple mass spectrometer (5500 QTRAP, AB SCIEX) in the multiple reaction monitoring (MRM) mode. Metabolites were detected in electrospray negative-ionization and positive-ionization modes. Samples (2 μL) were injected sequentially. The Waters ACQUITY UPLC HSS T3 C18 (1.8 μm, 2.1 mm 100 mm, Wasters)was heated to 40°C at a flow rate of 400μL/min. The gradient was started with the solvent and water/acetonitrile ratio at 95%/5%(V/V) for 11 min, after which the volume ratio was changed to 5%/95% for 1 min over 11 min, and finally to 95%/5% at 12.1 min, lasting 15 min.

#### Data processing and analysis

The local database was used to analyze the metabolite information, and quantitative information was determined in the MRM mode. The identified substances were analyzed by Kyoto Encyclopedia of Genes and Genomes (KEGG), and the differences among different samples were analyzed by principal component analysis (PCA) and Orthogonal partial least squares discriminant analysis (OPLS-DA).

### Part 3: Effects of different metabolites on *O. communa*

Based on the results of metabolomics, we selected the significantly upregulated secondary metabolites of common ragweed exposed to heat waves and high concentrations of CO_2_; then, we set up a bioassay experiment to detect the effect of upregulated secondary metabolites on leaf beetle survivorship.

#### Bioassay test

Pure products of the purchased substances were prepared into 2000 ppm, 1500 ppm, 1000 ppm, and 500 ppm series multiples with acetone (analytical purity) for reserved use. The fresh common ragweed leaves were cut into discs with a diameter of approximately 1–2 cm. The leaf discs were put into the doubling solutions of three substances for 5 s and then removed. After the solvent evaporated, the leaf discs were placed in a Petri dish (diameter 9 cm) padded with filter paper. The bottom layer of the filter paper was put into a piece of absorbent cosmetic cotton to moisturize the leaf discs. After starvation treatment of the 2^nd^ instar larvae for 8 h, they were placed on the leaves in the Petri dish; fifteen test larvae were treated, with four repetitions, and acetone was set up as a blank control. The 1°C, night temperature of 25± 70±5%, illumination condition of 14:10 L:D, light intensity of 30,000 lux). The larvae were fed for 5 days with leaves soaked in metabolite doubling solution. After 5 days, the common ragweed leaves were replaced with fresh leaves until pupation of the larvae. Fresh leaves were replaced every 24 h. The larvae were investigated, and their daily mortality in each period was recorded.

#### Statistical analysis

Data were checked for normality as appropriate (*P*>0.05) and, if needed, were arcsine square-root or log-transformed. The data for growth, foliar chemistry, and the biological parameters of the *O. communa* adults fed on common ragweed under the four treatments were analyzed using a two-way analysis of variance (ANOVA) via IBM SPSS Statistics 20; the differences between means were analyzed by one-way ANOVA via IBM SPSS Statistics 20 (LSD test, P¼ 0.05; SPPS, Inc., Chicago, IL, USA).

Life table parameters were estimated using the TWO-SEX-MS Chart program (Chi, 2012) based on the age-stage and two-sex life table theory (Chi, 1988). The age-stage-specific survival rate (Sxj) (x¼age, j¼stage), age-stage-specific fecundity (fxj), age-specific survival rate (lx), age-specific fecundity (mx), and population parameters (*r*, the intrinsic rate of increase; k, the finite rate of increase; R0, the net reproductive) were analyzed.

For the metabolomics data, the SIMCAP 14 software (Umetrics, Umeå, Sweden) was used for all multivariate data analyses and modeling. Data were mean-centered using Pareto scaling. Models were built on PCA. The discriminating metabolites were obtained using a statistically significant threshold of fold change (FC) and two-tailed Student’s t-test (*p* value) on the normalized raw data. The *p* value was calculated by one-way ANOVA for multiple groups. Metabolites with FC greater than 1.5 and a*p* value less than 0.05 were considered statistically significant metabolites. FC was calculated as the logarithm of the average mass response (area) ratio between two arbitrary classes. The identified differential metabolites were used to perform cluster analyses using the R package.

## Acknowledgements

This work was supported by the National Natural Science Foundation of China (31972340 and 31672089).

## Author contributions

ZT and CM contributed equally to this work. ZZ designed the study and revised the manuscript. Together, ZYT and CM performed the research and wrote the manuscript. CZ performed the data analyses. YZ, XG, and ZQT participated in sample collection and data sorting. HC and JG provided guidance during the data analysis.

## Abbreviations

ANOVA: Analysis of variance
BapNA: Benzoyl-L-arg-p-nitroanilide FC Fold change
KEGG: Kyoto Encyclopedia of Genes and Genomes MRM Multiple reaction monitoring
PCA: Principal component analysis PPFD Photosynthetic photon flux density
QC: Quality control
RH: Relative humidity

---

**Supplementary Fig. 1.**
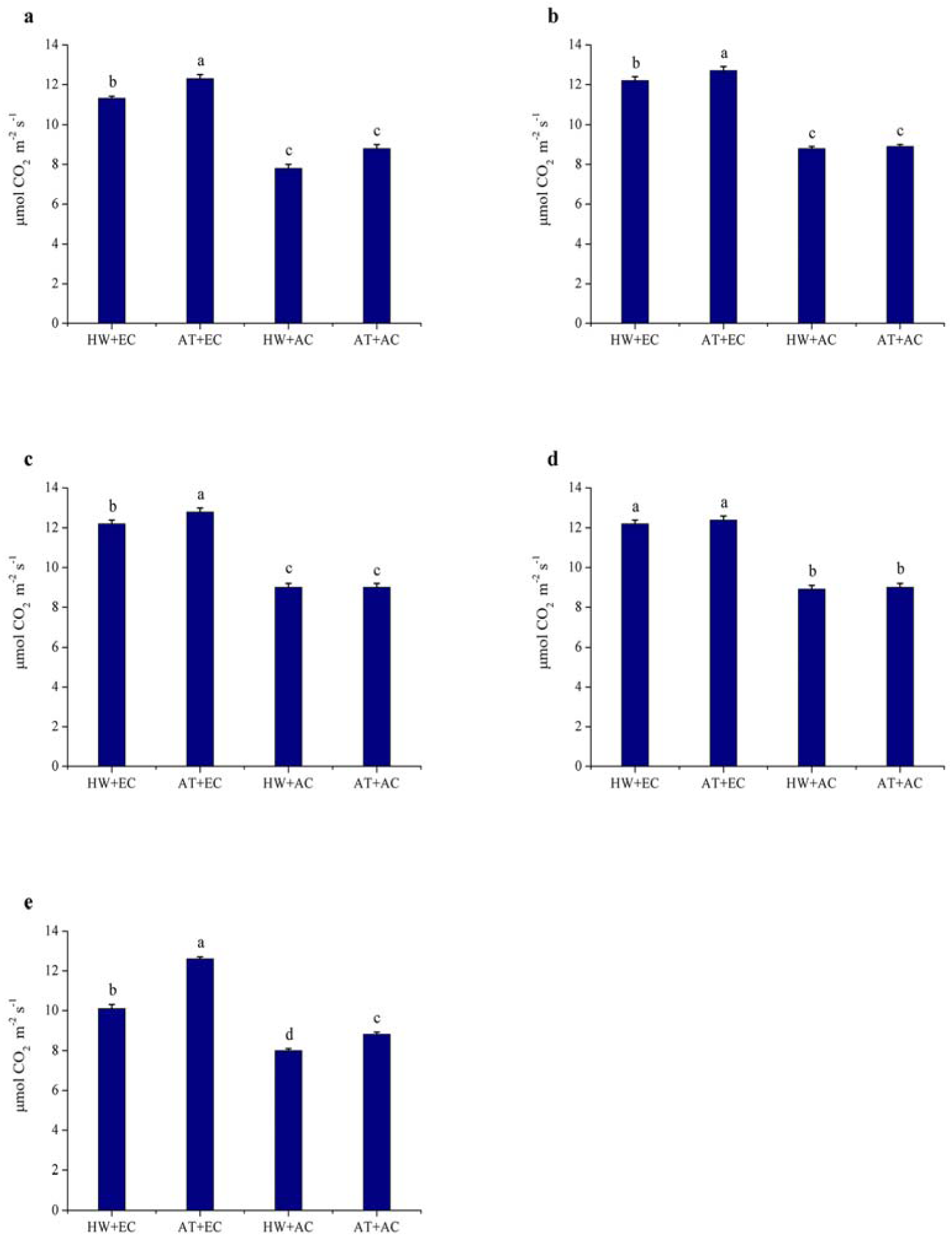
Leaf photosynthesis intensity (± SE) of *Ambrosia artemisiifolia* under different stress conditions. Data represented by columns bearing the same letters were not significantly different (LSD, *P* = 0.05). A, B, C, D, and E represent the 1^st^, 2^nd^, 3^rd^, 4^th^, and 5^th^ measurements, respectively. AT denotes the normal temperature condition; HW denotes the heat wave condition. EC and AC represent elevated atmosphere CO_2_ concentration and ambient atmosphere CO_2_ concentration, respectively.

**Supplementary Fig. 2.**
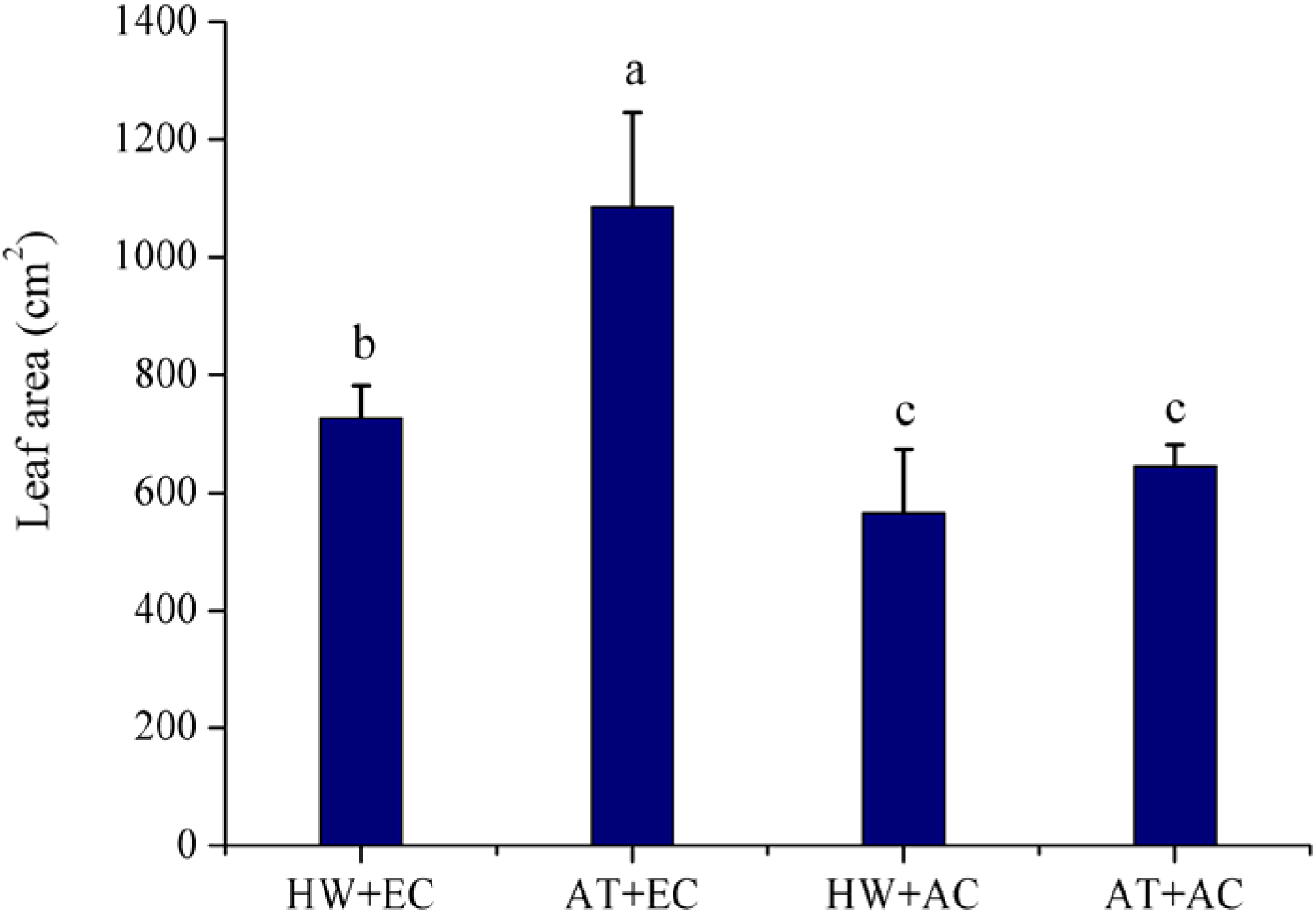
Total leaf area (±SE) of *Ambrosia artemisiifolia* under different stress conditions. Data represented by columns bearing the same letters were not significantly different (LSD, *P* = 0.05). AT denotes the normal temperature condition; HW denotes the heat wave condition. EC and AC represent elevated atmosphere CO_2_ concentration and ambient atmosphere CO_2_ concentration, respectively.

**Supplementary Fig. 3.**
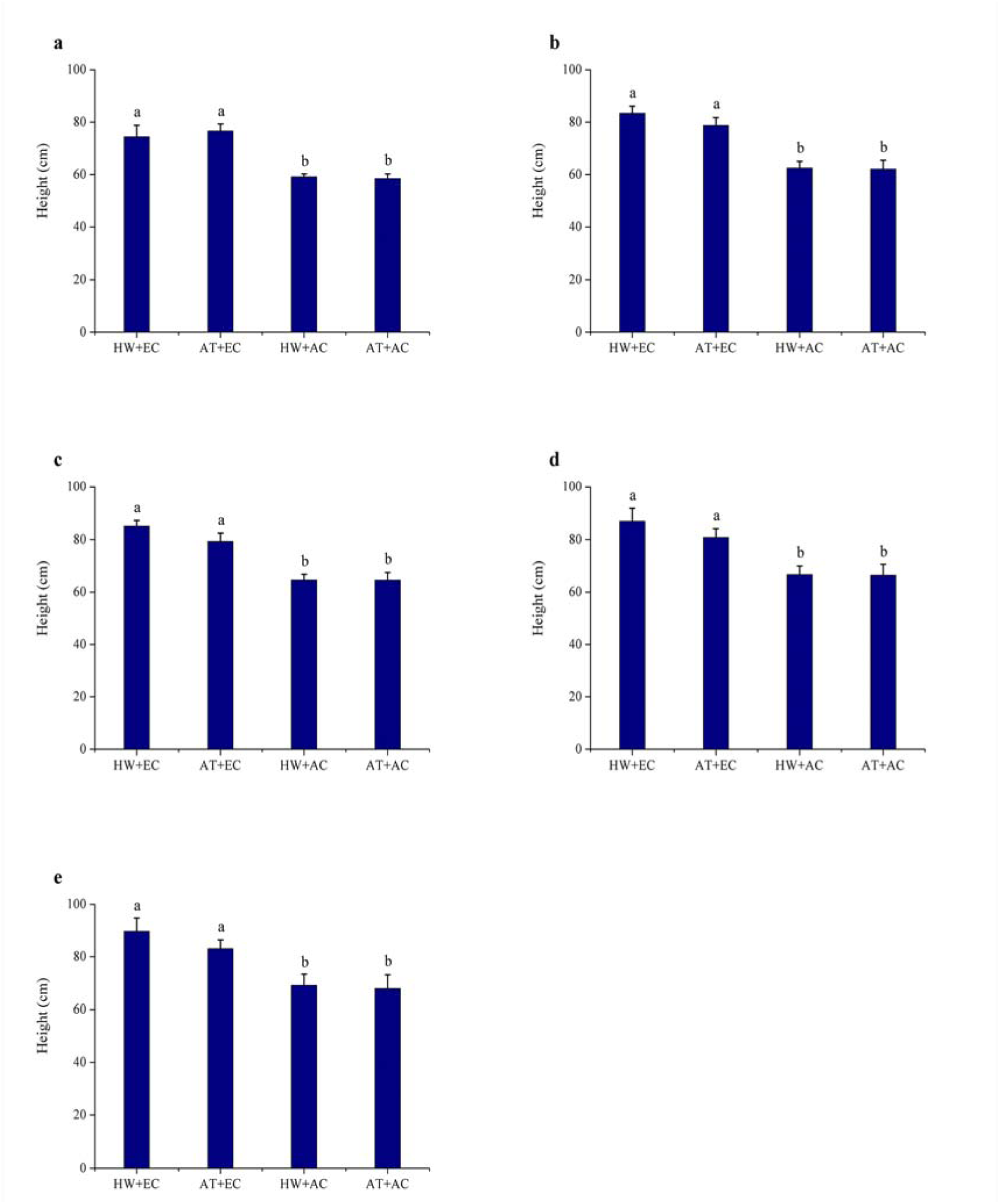
Plant height (±SE) of *Ambrosia artemisiifolia*under different stress conditions. Data represented by columns bearing the same letters were not significantly different (LSD, *P* = 0.05). A, B, C, D, and E represent the 1^st^, 2^nd^, 3^rd^, 4^th^, and 5^th^ measurements, respectively. AT denotes the normal temperature condition; HW denotes the heat wave condition. EC and AC represent elevated atmospheric CO_2_ concentration and ambient atmosphere CO_2_ concentration, respectively.

**Supplementary Fig. 4.**
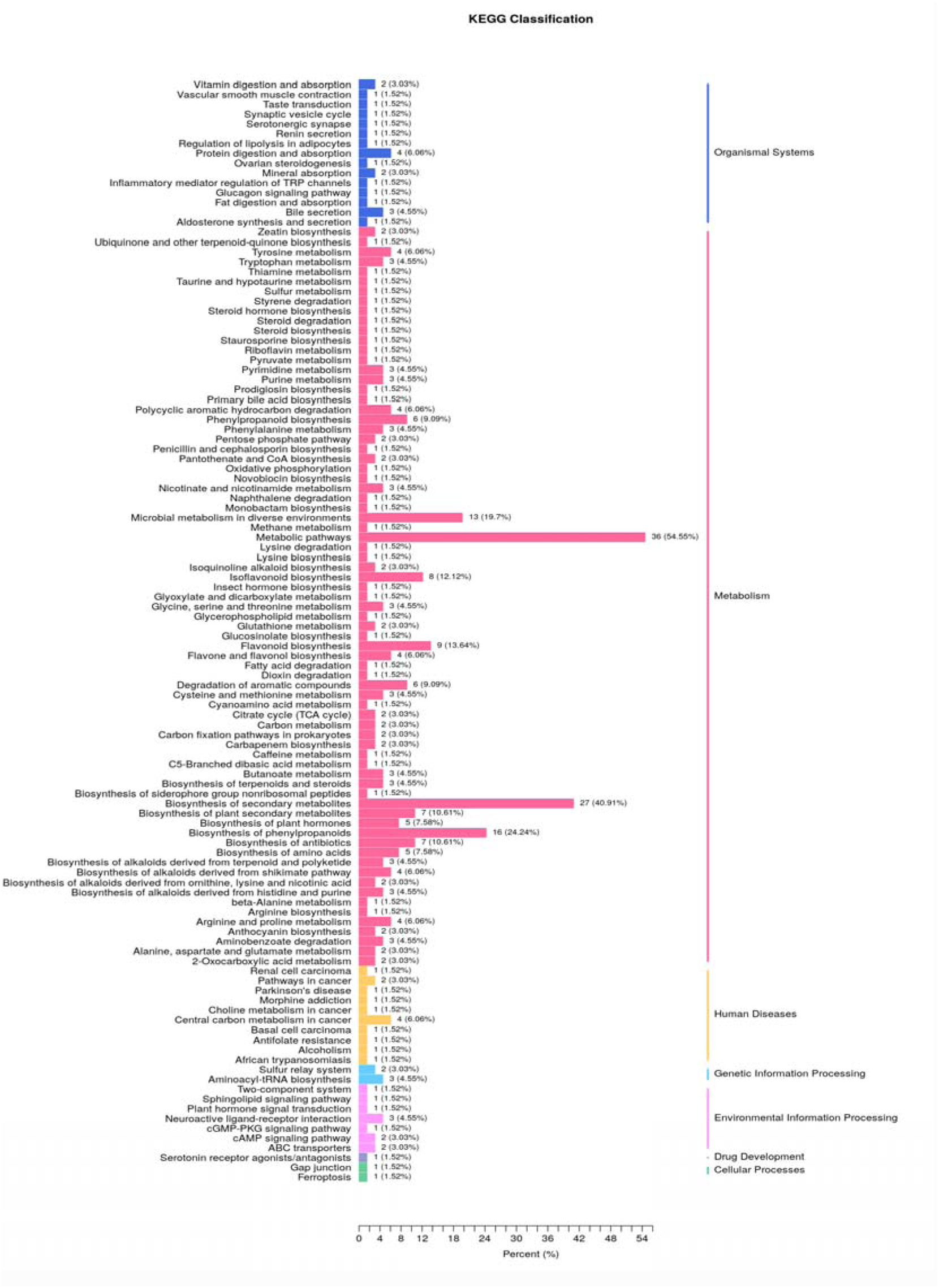
KEGG analysis of different metabolites between leaves of plants under AC+AT and EC+HW conditions.

